# Estimation of Species Abundance Based on the Number of Segregating Sites using Environmental DNA (eDNA)

**DOI:** 10.1101/2024.04.15.589638

**Authors:** Qiaoyun Ai, Hao Yuan, Ying Wang, Chenhong Li

**Author notes:** Department of Biochemistry and Molecular Biology, Michigan State University, East Lansing, MI 48823, USA. Genetics and Genome Sciences Program, Michigan State University, East Lansing, MI 48823, USA, Ecology. Evolution and Behavior Program, Michigan State University, East Lansing, MI 48823, USA. **Correspondence to:** Chenhong Li.

## Abstract

The advance of environmental DNA (eDNA) has enabled rapid and non-invasive species detection in aquatic environments. Although most studies focus on species detections, some recent studies explored the potential of using eDNA concentration to quantify species abundance. However, the differential individual DNA contribution to eDNA samples could easily obscure the concentration-species abundance relationship. We propose using the number of segregating sites as a proxy for estimating species abundance. Since segregating sites reflects the genetic diversity of the population, which is less sensitive to differential individual DNA contribution than eDNA concentration. We examined the relationship between the number of segregating sites and species abundance in silico, in vitro, and in situ using two brackish goby species, *Acanthogobius hasta* and *Tridentiger bifasciatus*. Analyses of the simulated data and in vitro data with DNA mixed from a known number of individuals showed a strong correlation between the number of segregating sites and species abundance (R^2^ > 0.9; P < 0.01). Results from the in situ experiment further validated the correlation (R^2^ = 0.70, P < 0.01), and such correlation was not affected by biotic factors, including body size and feeding behavior (P > 0.05). Results of the cross-validation test also showed that the number of segregating sites predicted species abundance with less bias and variability than the eDNA concentration. Overall, the number of segregating sites correlates stronger with species abundance and also provides a better estimate than eDNA concentration. This advancement can significantly enhance the quantitative capabilities of eDNA technology.

## INTRODUCTION

The environmental DNA (eDNA) technique has emerged as a prevalent approach in ecological studies due to its cost-effectiveness, enhanced efficiency, and non-invasive nature compared to traditional approaches (Furlan and Gleeson 2016; Hosler 2017; Sigsgaard et al. 2015). By sampling environment samples from various ecosystems, such as tropical freshwater (Robson et al. 2016), intermittent river (Bylemans et al. 2016), brackish water (Forsström and Vasemägi 2016), kelp forest (Port et al. 2016), and even terrestrial habitat (Bohmann and Lynggaard 2023), the eDNA technique can capture ambient DNA released by organisms, which is subsequently analyzed to detect species. The eDNA approach has been widely applied in biomonitoring, including endangered (Fukumoto, Ushimaru, and Minamoto 2015) and invasive species (Uchii, Doi, and Minamoto 2016), allowing for monitoring single or multiple species simultaneously (DiBattista et al. 2017). Additionally, the eDNA technique has also been expanded into population studies, which provide a practical and nondestructive approach to determining species composition (Madduppa et al. 2021), symbiotic relationships (Shinzato et al. 2018), complex species structures (DiBattista et al. 2017) and breeding habits (Sakata et al. 2017).

Another promising frontier in eDNA research is the estimation of species abundance. This application, while challenging, holds significant value for conservation ecology and population genetics. eDNA offers a non-invasive method to monitor changes in species abundance, particularly for invasive species, thereby informing more effective management strategies (Matthews and Whittaker 2015). Furthermore, species abundance is a key parameter in understanding population dynamics. Fluctuations in abundance are closely related to changes in various population attributes, including genetic diversity (Hughes et al. 2008) and selection pressures (Lankau and Strauss 2011). Several studies have explored the potential of using eDNA concentration as a proxy by relating it to species abundance (Chambert et al. 2018; Dunn et al. 2017:201; Hänfling et al. 2016; Yusishen et al. 2020). Some studies reported strong positive relationships between eDNA concentration and species abundance (Karlsson et al. 2022; Yates, Fraser, and Derry 2019), while others showed non-significant relationships (Brys et al. 2021). Those contradictory findings likely resulted from the influences of various biotic and abiotic factors on eDNA dynamics - shedding, decaying, and transporting, which collectively affect eDNA concentration (Barnes and Turner 2016). For example, Biological characteristics such as species (Ficetola et al. 2008; Jerde et al. 2011), tissue source (Merkes et al. 2014), life stage (Maruyama et al. 2014), and weight (Mizumoto et al. 2018) could affect the shedding rate of eDNA. Environmental conditions including temperature (Strickler, Fremier, and Goldberg 2015), UV intensity (Strickler et al. 2015), and microbial community (Poté, Ackermann, and Wildi 2009) play a role in determining the decaying rate of eDNA. Additionally, physical processes like water flow (Foppen et al. 2011) and precipitation/resuspension (Turner, Uy, and Everhart 2015) contribute to the transport of eDNA in the ecosystem. These factors combined lead to the uneven contribution of DNA from individuals among the population. Some individuals significantly influence the measured concentration, potentially leading to overestimation of their numbers, while others contribute less to the overall eDNA concentration and potentially are underrepresented in species abundance estimates. Thus, the differential contribution of individuals to the collected eDNA samples skews the species abundance estimates from eDNA concentration (Andres et al. 2023). Consequently, developing a metric resilient to differential individual contribution to eDNA samples is imperative for achieving an accurate and precise estimate of species abundance.

To establish a resilient metric against the differential individual contribution to eDNA samples, we propose using the number of segregating sites - polymorphic sites within the population - as a proxy for species abundance. As an indicator of genetic variation, the number of segregating sites rises with increasing sample size in equilibrium populations (Tajima 1989; Takahashi and Tajima 2017). The inherent redundancy nature of segregating sites ensures that each individual is accessed only once regardless of the number of copies of DNA detected in the samples. We investigated the relationship between the number of segregating sites and species abundance through a comprehensive, three-tiered experimental design. Our study began with an in silico analysis, where we tested the relationship using completely simulated data generated at various mutation rates (Fig. 1a). The in silico study was then complemented by an in vitro analysis, in which we simulated eDNA samples by subsampling sequences from tissues of two brackish water goby species, *Acanthogobius hasta* and *Tridentiger bifasciatus* (Fig. 1b). Finally, to evaluate our findings in a more natural context, we conducted in situ (mesocosm) experiments to analyze eDNA collected from aquariums (Fig. 1c). Results revealed that the number of segregating sites extracted from enriched sequences showed a stronger correlation and a better estimate of species abundance than eDNA concentration.

**Fig. 1.**
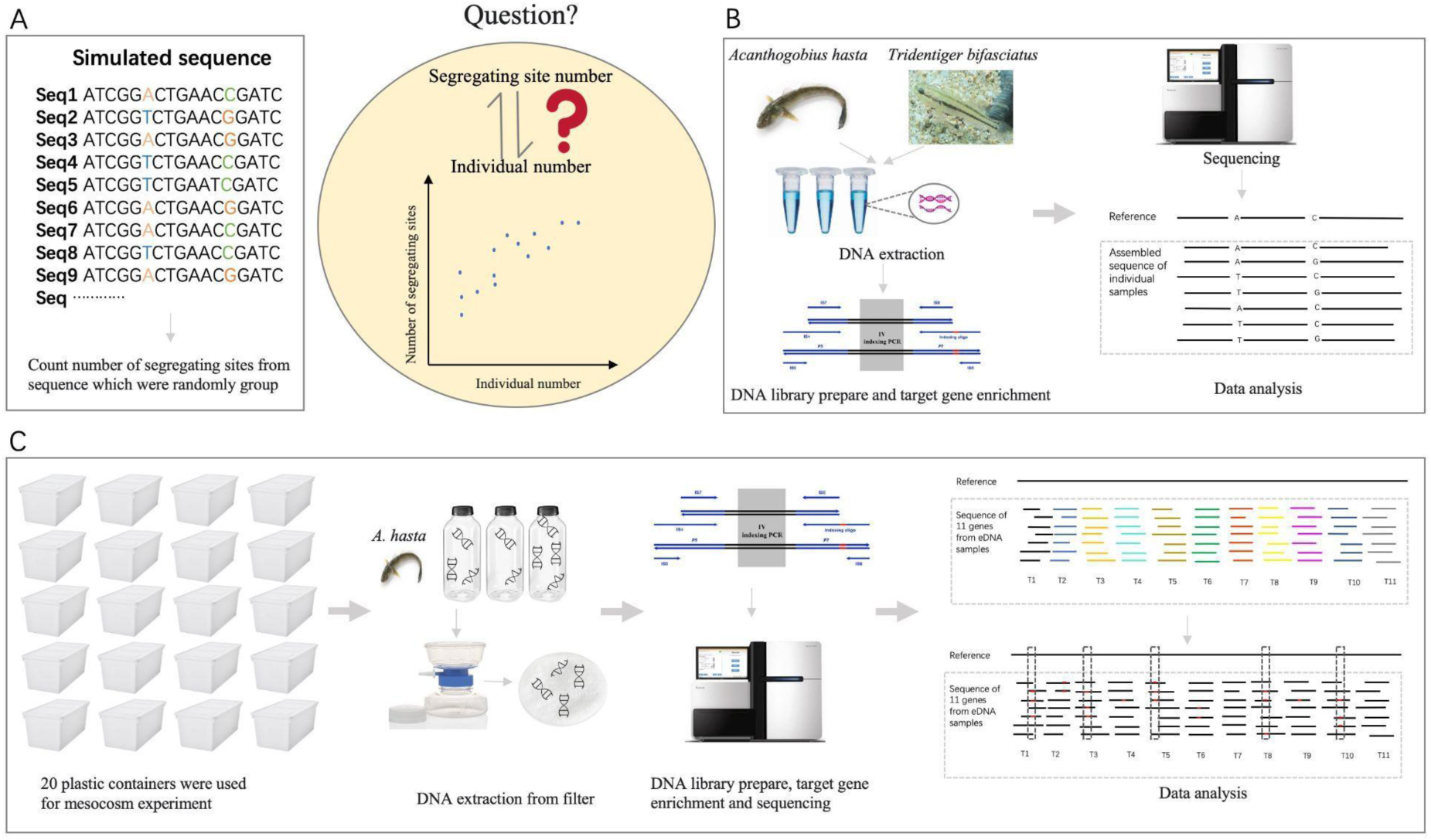
Three main components of this study. (A) In silico study. We tested the correlation between the number of segregating sites and the species abundance using simulated data. (B) In vitro study. We designed markers and synthesized baits to capture target genes from tissues of *A. hasta* and *T. bifasciatus* individuals. Datasets mixed from captured sequences were used to assess the correlation between the number of segregating sites and the species abundance. (C) In situ (mesocosm) study. We conducted controlled mesocosm experiments using *A. hasta*. Segregating sites were extracted from captured sequences in eDNA samples, and used to evaluate their correlation to species abundance and their performance in estimating species abundance.

## 2 MATERIAL AND METHODS

### 2.1 In Silico Experiment

We first assessed the relationship between the number of segregating sites and species abundance by entirely simulated sequences. The length of simulated sequences was set at 17,000 bp. The number of simulated sequences/individuals was 1,000, and sequences were generated at the mutation rate of 10^-6^ /bp/gen. To account for mutation rate variation among different species, we also generated another two datasets at the mutation rate of 10^-7^ /bp/gen and 10^-8^ /bp/gen. These mutation rates represent a biologically meaningful range for vertebrates (Allio et al. 2017). All data were generated using the software Fastsimcoal2 (Excoffier and Foll 2011). A subset of sequences were randomly chosen from the simulated data, ranging from 20 to 980 sequences with intervals of 20. Selected sequences were aligned using MUSCLE v1.0 (Edgar 2004), and then the number of segregating sites was counted from alignments. The simulation process was repeated three times at each specified number of sequences. The correlation between the number of segregating sites and the number of individuals/sequences was estimated by regression analysis using Microsoft Excel, in which the number of segregating sites was the dependent variable (y), the number of individuals/sequences was the independent variable (x) and the significance of the correlation was estimated using R^2^ and p-value. The simulated sequences are available in Dryad at https://doi.org/10.5061/dryad.w3r2280zz.

### 2.2 Target selection

In our study, we employ target enrichment rather than metabarcoding to collect DNA from eDNA samples. Target enrichment offers superior coverage to PCR, allowing us to extract more segregating sites from wider genomic regions for species abundance estimates (Li et al. 2023). Our goal is to design a single set of RNA baits for enrichment that can be applied to a wide range of species. To achieve this versatility, we focus on identifying mtDNA regions that are highly conserved across diverse species. This strategy ensures better enrichment across multiple species while utilizing variable flanking regions to extract segregating sites.

We conducted a comprehensive alignment of mitochondrial genome sequences from a broad spectrum of vertebrates, including *Danio rerio*, *Cyprinus carpio*, *Oryzias latipes*, *Tetraodon nigroviridis*, *Gasterosteus aculeatus*, *Lepisosteus oculatus*, *Anguilla japonica*, *Gadus morhua*, *Oreochromis niloticus*, *Homo sapiens*, *Xenopus tropicalis*, *Anolis carolinensis*, *Callorhinchus milii*, and *Gallus gallus* (Table S1). We selected regions with an average pairwise similarity above 0.8 and a minimum length of 120 bp to ensure sufficient conservation. This rigorous selection process yielded 11 segments from the mitochondrial genome that met our criteria. These segments were subsequently employed as target markers for both our in vitro and in situ experiments.

### 2.3 In Vitro Experiment

Before using 11 markers to enrich targets from eDNA samples, we need to test the performance of using those markers to enrich DNA extracted from tissue samples. Thus, we examined the correlation between the number of segregating sites and species abundance using enriched sequences to get a sense of correlation from actual populations. We selected two species of fish for the in vitro experiment: *A. hasta* (92 samples) were bought from fishermen who caught fish from the East China Sea in August 2016, and *T. bifasciatus* (78 samples) caught from Dishui Lake in Shanghai. The DNA of two species was extracted from fin clips using an Ezup Column Animal Genomic DNA Purification Kit (Sangon, Shanghai, China). The purified DNA was quantified with a NanoDrop 3300 Fluorospectrometer (Thermo Fisher Scientific, Wilmington, DE, U.S.A.) and visualized by agarose gel electrophoresis. The primers were designed according to the mitochondrial genome sequence of *A. hasta* (NCBI accession: AY486321) and *T. trigonocephalus* (NCBI accession: NC_029738) (Table S1). We evaluated the specificity of the primers using the software Primer3.

RNA baits used for target enrichment were RT-PCR products of the 11 target segments. TakaRa Ex Taq® (RR001 Takara) was used to amplify 11 target DNA segments using the DNA from an *A. hasta* sample (CL1176_43) as a template. Eleven pairs of primers used to amplify target segments are listed in Table S4. The PCR products were checked using agarose gel electrophoresis and mixed as template transcription. Then, we followed the manual of MEGAshortscript™ kit (Thermo Fisher Scientific, Wilmington, DE, U.S.A.) to transcribe PCR products to RNA transcripts. The concentration of purified transcribed product was measured and then stored at -80 °C until the gene enrichment.

The process of gene enrichment followed Li et al.’s protocol (Li et al. 2013). We sheared the 1,000 ng purified DNA into around 250 bp using a Covaris M220 Focused-ultrasonicator (Woburn, Massachusetts, U.S.A.). Sheared DNA was used to prepare libraries with two different indexes incorporated in both ends. All indexed libraries were mixed into one tube equimolarly for gene enrichment and Illumina sequencing on a HiSeq 2500 sequencer (Illumina, Inc, San Diego, CA, U.S.A.).

The enriched sequences were mapped to mtDNA reference sequences of *A. hasta* and *T. bifasciatus* to get potential target sequences, which were further assembled using the software Trinity2.06 (Grabherr et al. 2011). We evaluated the enrichment quality based on raw read numbers, mapped read numbers, enriched loci numbers, and sequencing depth.

We constructed a phylogenetic tree using the assembled sequences to validate the species identity of our collected samples. Assembled sequences of 92 *A. hasta* and 78 *T. bifasciatus* were used to build maximum likelihood trees using GTRGAMMA model in RAxML v8.0.0 (Stamatakis 2014) with *Mugilogobius abei* (NCBI accession: NC_023353.1) as outgroup. We used 92 sequences enriched from *A. hasta* and 78 sequences enriched from *T. bifasciatus* to evaluate the correlation between the number of segregating sites and the number of individuals.

The sequences were randomly selected across diverse group sizes, ranging from 5 to 90 for *A. hasta*, and 5 to 70 for *T. bifasciatus*. We generated three repeats for each group size. The sequences in each group were aligned using MUSCLE v1.0 (Edgar 2004), then the number of segregating sites was counted using MEGA7 (Kumar, Stecher, and Tamura 2016). The correlation between the number of segregating sites and the number of individuals was evaluated by regression analysis using Microsoft Excel. The number of segregating sites was set as the dependent variable (y), while the number of individuals was the independent variable (x) and the significance of the correlation was estimated by R^2^ and p-value.

### 2.3 In Situ Experiment (Mesocosm experiment)

Finally, we conducted a mesocosm experiment to evaluate the relationship between the number segregating sites and species abundance in water samples. *A. hasta* used for the mesocosm experiment were captured from the East China Sea in September 2017. The fish were kept in dechlorinated water with 5‰ salinity and fed with frozen shrimp before the experiment. The water in the aquaria was changed at least four times, and the dead fish were removed in a timely manner until all the fish were accustomed to the aquaria. All equipment, including containers, air stones, and pipes were sterilized one week before the experiment. To reduce the mortality of fish, we did not weigh the individuals for size grouping, but divided them into four groups according to the body length (< 10 cm, 10-12.5 cm, 12.5-15 cm, > 15 cm). The average weight of fish was measured after the experiment (Table 1). Twenty 65 L plastic containers were prepared to keep *A. hasta* (Table 1), and one aquarium without fish was set as negative control. We set five fish number gradients, including 7, 14, 21, 28, 35. To account for the effect of feeding behavior in the estimation of species abundance, *Penaeus vannamei* was cut into pieces to feed the fish, at approximately 5% of the estimated total weight of fish in each aquarium. One liter of water in every aquarium was collected after three days. The collected water was filtered using 47-mm GF/F glass filters (pore size c. o.7 μm; Whatman International, Ltd), then the filters were soaked in pure ethanol and stored at -80 °C until DNA extraction. DNA samples were extracted using an Ezup Column Animal Genomic DNA Purification Kit (Sangon, Shanghai, China). The purified eDNA was checked using agarose gel electrophoresis and stored at -20 °C until the target gene enrichment and qPCR.

**Table 1.** Design of the mesocomic experiments.

| Feeding\No. fish | 7<sup>a</sup> | 14 | 21 | 28 | 35 |
| --- | --- | --- | --- | --- | --- |
| F<sup>b</sup> | W<sub>1</sub>(6)<sup>c</sup> | W<sub>2</sub>(12) | W<sub>3</sub>(18) | W<sub>4</sub>(39) | W<sub>1</sub>(11) |
| F<sub>0</sub> | W<sub>2</sub>(13) | W<sub>1</sub>(7) | W<sub>4</sub>(38) | W<sub>3</sub>(24) | W<sub>2</sub>(15) |
| F | W<sub>3</sub>(19) | W<sub>4</sub>(44) | W<sub>1</sub>(7) | W<sub>2</sub>(13) | W<sub>3</sub>(20) |
| F<sub>0</sub> | W<sub>4</sub>(49) | W<sub>3</sub>(16) | W<sub>2</sub>(14) | W<sub>1</sub>(8) | W<sub>4</sub>(42) |

<sup>a</sup>The number (7, 14, 21, 28 and 35) indicates the number of fish kept in each aquarium.
<sup>b</sup>F means the fish were fed during the experiment. F<sub>0</sub> means the fish were not fed.
<sup>c</sup>The number in parentheses is the average weight of fish in this aquarium. The number in
subscript denotes the rank of the average weight of fish in the treatment group. Smaller rank in
subscript means lower average fish weight in this aquarium.

For better target enrichment efficiency, we redesigned primers for the in situ experiment according to sequences we obtained from the in vitro experiment (Table S4). PCR products were further transcribed and prepared as baits following instructions in section 2.3. By checking agarose gel electrophoresis, we found that the maximum length of purified eDNA was too large for next-generation sequencing. Thus, we sheared DNA to around 250 bp using a Covaris M220 Focused-ultrasonicator (Woburn, Massachusetts, U.S.A.). Libraries preparation, target gene enrichment and sequencing followed the above procedure. After preprocessing raw reads including trimming adaptors, removing low-quantity reads, and PCR duplicates, the reads of each sample were mapped to the reference of *A. hasta* using BWA-MEM (Li and Durbin 2010). The number of segregating sites was counted from the targeted regions as well as flanking sequences, spanning 50 bp upstream and downstream of the targeted regions. We removed polymorphic sites with sequencing depth below seven and minor allele frequencies below 3% to filter out potentially erroneous segregating sites stemming from PCR stutter and sequencing errors (Fumagalli 2013).

We also accessed the eDNA concentration of each sample. Among the 11 target segments, T2, the second segment, was chosen as the target sequence for qPCR due to its length (140 bp) falling within the recommended range for qPCR amplicons (Van Holm et al. 2021). To create standard samples, we amplified the T2 target region by TakaRa Ex Taq® (RR001 Takara) using the genomic DNA of CL1176_43 as the template. The amplified product was purified by gel extraction using DNA gel extraction kit (Qiagen, Hilden). The purified product was further diluted to create six standard samples for qPCR, with concentrations ranging from 1.75 ng/L to 1.75×10^6^ ng/L. The qPCR procedure followed the protocol outlined in KAPA SYBR FAST Universal qPCR Kit (KAPABIOSYSTEMS, U.S.A). We assessed the specificity of the eDNA amplified products through a melting curve analysis using the online BIO-RAD melting curve generator. The copy number was estimated using the online BIO-RAD copy number calculator. The number of DNA copies was estimated as the mean value of three repeats for the same water sample (Fig. S2). We averaged the number of DNA copies across replicates and converted this value to eDNA concentration (copies/L).

Statistical analysis was performed using Stata (https://www.stata.com/). Fixed effect models were used to determine if fish number, body size, and feeding behavior affected the number of segregating sites or eDNA concentration. We also constructed another simpler fix effect model, which only considered the fish number as the variable. We used the F-test, Akaike Information Criterion (AIC), and Bayesian information criterion (BIC) to discern whether species abundance is the only factor attributed to the impact on the number of segregating sites or eDNA concentration.

To take a step further from correlation analysis, we conducted an exhaustive leave-*p*-out cross-validation test to compare the performance of estimating species abundance using segregating sites and eDNA. Specifically, *p* samples were randomly selected from the mesocosm dataset as testing samples, while the rest served as training samples. *p* ranges from 1 to 10. Training samples were used to establish regression models between the number of segregating sites/eDNA concentration and species abundance. Then, those models were applied to estimate the species abundance in test samples. Notably, for every value of *p*, we exhaustively accessed the performance for every possible combination of samples. If *p* was greater than one, the species abundance of the same sample could be estimated multiple times by models built based on different training samples. We evaluated estimation performance using absolute bias, i.e., the absolute difference between ground truth and estimated species abundance. We further averaged absolute bias for the same sample when assessing performance. The precision of the estimation of each sample was calculated using the standard deviation of predicted species abundance.

## 3 RESULTS

### 3.1 In Silico Analysis

Simulation analysis revealed a strong positive correlation between the number of segregating sites and species abundance across all mutation rates. All regression models had R^2^ above 0.9. The regression coefficient for the number of segregating sites increased with mutation rate, reflecting the increased occurrence of segregating sites in the mtDNA of species with higher mutation rates (Fig. 2). Regression models fit better with log-transformed species abundance rather than its absolute value, since the number of segregating sites reached saturation when sampling enough of the number of individuals.

**Fig. 2.**
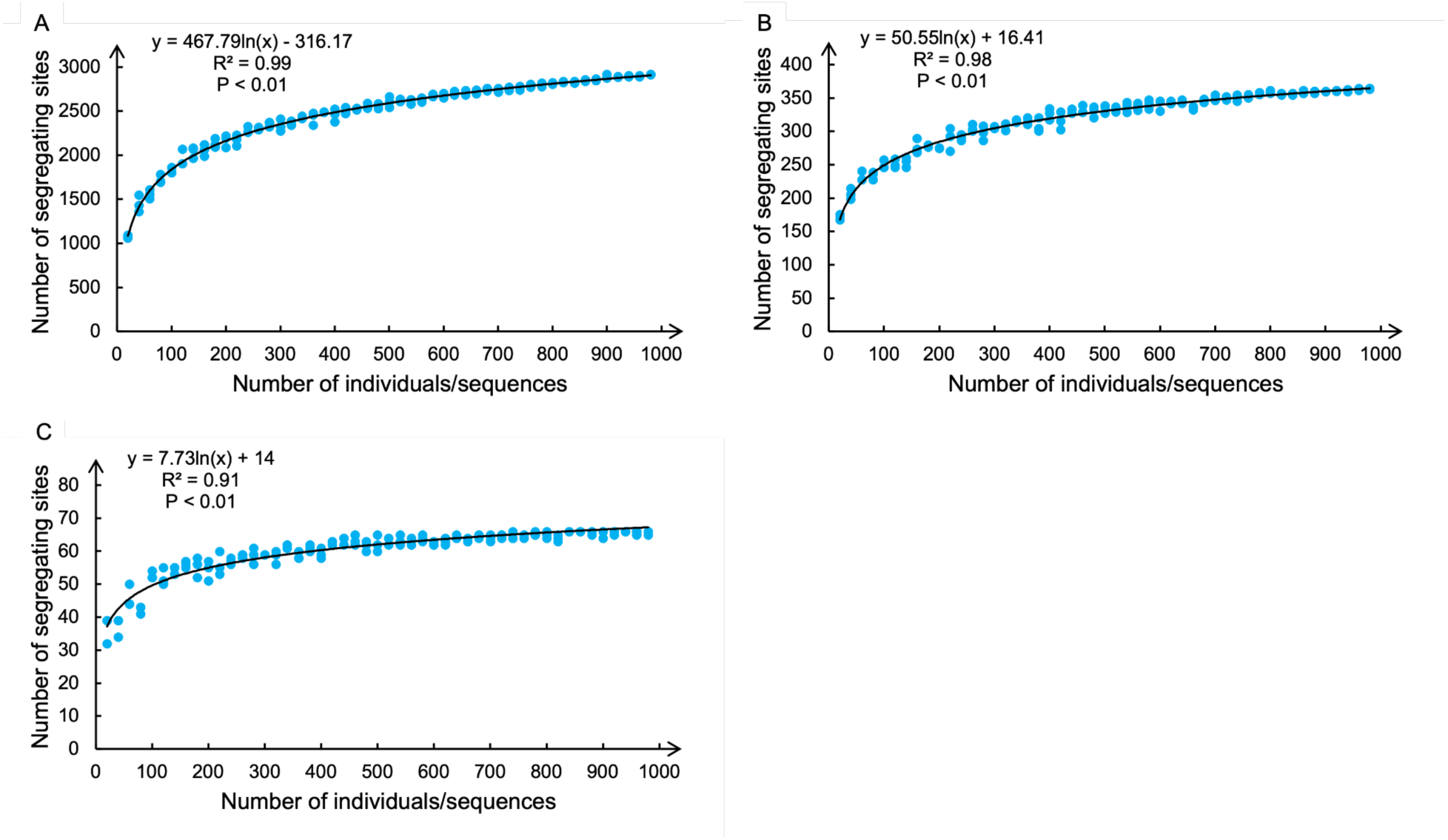
Relationship between the number of segregating sites and the number of simulated individuals/sequences from in-silico analysis. Sequences of 1,000 individuals, each with a length of 17,000 bp, were simulated with a mutation rate of 10^-6^ /bp/gen (A), 10^-7^ /bp/gen (B) and 10^-8^ /bp/gen (C) respectively. For each mutation rate, 20 to 980 samples were randomly drawn from the population, and the number of segregating sites was counted. The sampling process was repeated three times for each number of individuals. Each dot in the plots represents one random sampling. The fitted line represents the relationship between the number of segregating sites and the species abundance.

### 3.2 In Vitro Analysis

Before the mesocosm experiment, we used baits designed from 11 target segments to enrich DNA from tissues of *A. hasta* and *T. bifasciatus*, including target segments and their flanking regions (Table S1). The result revealed the successful enrichment of the 11 targets in all samples. After quality check and PCR duplication removal, an average of 5,304,806 bp reads for *A. hasta* and 46,355 bp for *T. bifasciatus* remained. Among those, an average of 16,354 bp reads (9%) were mapped to the assembly of enriched sequences, with an average sequencing depth of 119 for *A. hasta* and 35 for *T. bifasciatus* (Table S2). The length of aligned sequences of *A. hasta* was 9,183 bp on average. For *T. bifasciatus*, the length of aligned sequences was 6,469 bp on average. We observed 507 segregating sites in *A. hasta*, and 163 segregating sites in *T. bifasciatus* respectively. The raw reads and the assembled sequences are available at NCBI Sequence Read Archive (SRA) with accession numbers PRJNA1100443.

Using *M. abei* as an outgroup (NCBI accession: NC_023353.1), phylogenetic analyses showed that samples were grouped into two clades. The sequences of *A. hasta* were clustered with the sequence of *A. hasta* retrieved from GenBank (NCBI accession: AY486321). The sequences of *T. bifasciatus* were grouped with *T. trigonocephalus* (NCBI accession: NC_029738) as their close sister species (Fig. S1). The topology of the phylogenetic tree and the grouping pattern of samples ensure the correct identification of the species.

We examined the relationship between the number of segregating sites and species abundance using enriched sequences from tissues. Regression analysis revealed that the number of segregating sites was positively correlated with the species abundance in both *A. hasta* and *T. bifasciatus* (Fig. 3). R^2^ values for regression models of both species exceeded 0.9. In contrast to the in silico study, the absolute value of species abundance fit better with models than log transformed form since the number of segregating sites observed in collected individuals was far from reaching saturation. A strong correlation was observed between the number of segregating sites and species abundance in the enrichment data from both *A. hasta* and *T. bifasciatus*. However, *A. hasta* exhibited higher genetic variability, resulting in stronger correlations. Consequently, only *A. hasta* was selected for the subsequent mesocosm experiments.

**Fig. 3.**
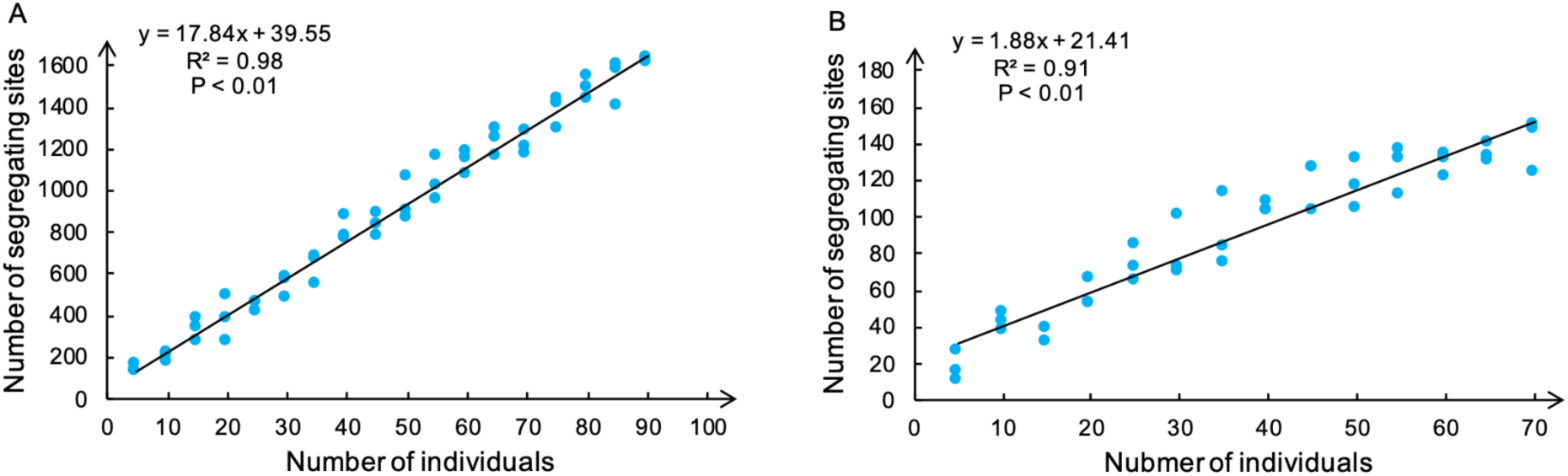
Relationship between the number of individuals and the number of segregating sites observed in the assembled sequences of *A. hasta* and *T. bifasciatus*. For *A. hasta*, 5 to 90 individuals were randomly sampled from the population, and the number of segregating sites was counted (A). Similarly, for *T. bifasciatus*, 5 to 70 were randomly sampled from the population, then the number of segregating sites was counted (B). The sampling process was repeated three times for each number of individuals. Each dot in the plots represents one random sampling. Fitted lines represent the relationship between the number of individuals and the species abundance.

### 3.3 In Situ (Mesocosm) Experiments

We enriched target sequences from eDNA samples using baits designed based on 11 targeted segments. After quality filtering and deduplication, 164,145,279 reads remained. Among those, 20,571 reads (0.59 %) were successfully mapped to the majority consensus assembly of *A. hasta* from in vitro analysis, with an average sequencing depth of 197 (Table S3). Subsequently, a depth filter was applied to remove potential false positive polymorphic sites with sequencing depth below seven and minor allele frequency below 3% (Fumagalli 2013). The raw reads are available at SRA with accession numbers PRJNA1100453.

Mesocosm Experiments revealed a strong and significant positive correlation between the number of segregating sites and species abundance (r = 4.09, p < 0.01). Although we also observed a positive relationship (r = 3.08, p < 0.01) between eDNA concentration and species abundance, species abundance accounted for a smaller proportion of variation in the eDNA concentration (R^2^ = 0.46) compared to the number of segregating sites (R^2^ = 0.70) (Fig. 4).

**Fig. 4.**
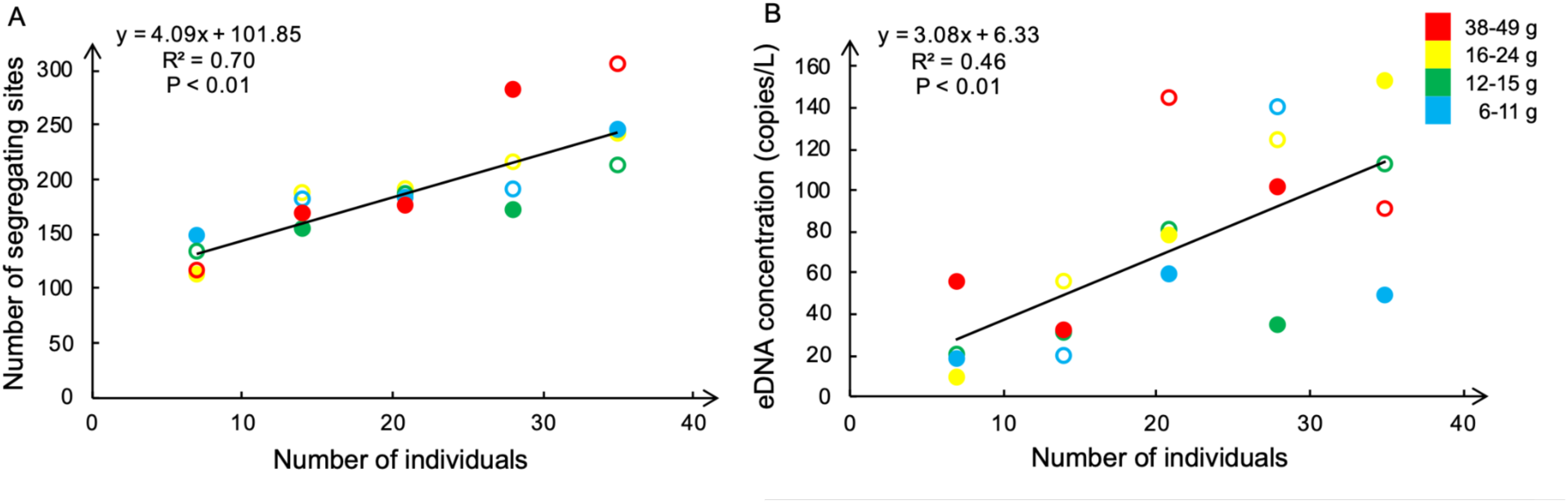
Relationship between the number of individuals and the number of segregating sites in eDNA samples (A), or between the number of individuals and the eDNA concentration (B). The eDNA concentration was estimated based on the number of DNA copies at locus T2 using qPCR. The colors of the dots depict the weight range of average weight for individuals in each aquarium: blue (6-11 g), green (12-15 g), yellow (16-24 g), and red (38-49 g). The solid dots indicate that the individuals in these aquariums were fed, and the circles indicate that the individuals were not fed. The number of individuals stocked in each tank ranged from 7,14, 21, 28 to 35. Four replicates were set up at each stocking density level, with different average weights and feeding treatments.

According to the F-test, AIC and BIC value, species abundance dominated the impact on the number of segregating sites. At the same time, mean weight and feeding behavior showed no significant influence (F-test p = 0.38). On the contrary, mean weight or feeding behavior demonstrated a nearly significant effect on eDNA concentration (F-test p = 0.07) (Table 2).

**Table 2.** Comparison of linear models for predicting the number of segregating sites or eDNA concentration using fish number, mean weight and feeding behavior.

| Variable | Segregating sites |  | eDNA concentration |  |
| --- | --- | --- | --- | --- |
|  | Model 1 | Model 2 | Model 1 | Model 2 |
| Fish number (Coef.) | 4.09** | 4.08** | 3.08** | 3.06** |
| Mean weight (Coef.) |  | 2.54 |  | -25.95 |
| Feeding (Coef.) |  | 0.63 |  | 0.66 |
| R<sup>2</sup> | 0.70 | 0.73 | 0.46 | 0.60 |
| AIC | 191.6 | 193.5 | 200.7 | 198.9 |
| BIC | 193.6 | 197.5 | 202.7 | 202.9 |
| F-test (p-value)<sup>a</sup> |  | 0.38 |  | 0.07 |

\* P < 0.05; \*\* P < 0.01.
<sup>a</sup> Comparing model 1 vs. model 2 using F-test statistics.

The cross-validation results indicated that the number of segregating sites is a more accurate and precise proxy for estimating species abundance. The median absolute bias of species abundance predicted based on the number of segregating sites was slightly lower than the one derived from eDNA concentration. This difference in median absolute bias became more pronounced when fewer samples were used as training data. Additionally, the upper limit of bias significantly decreased when predicting species abundance using the number of segregating sites compared to eDNA concentration (Fig. 5A). We evaluated the precision of species abundance estimation by examining the standard deviation of predictions for the same sample across different training datasets. As anticipated, we observed increasing standard deviation when fewer samples were served as training data. The standard deviation of species abundance estimates based on segregating sites was consistently lower than those predicted using eDNA concentration (Fig. 5B).

**Fig. 5.**
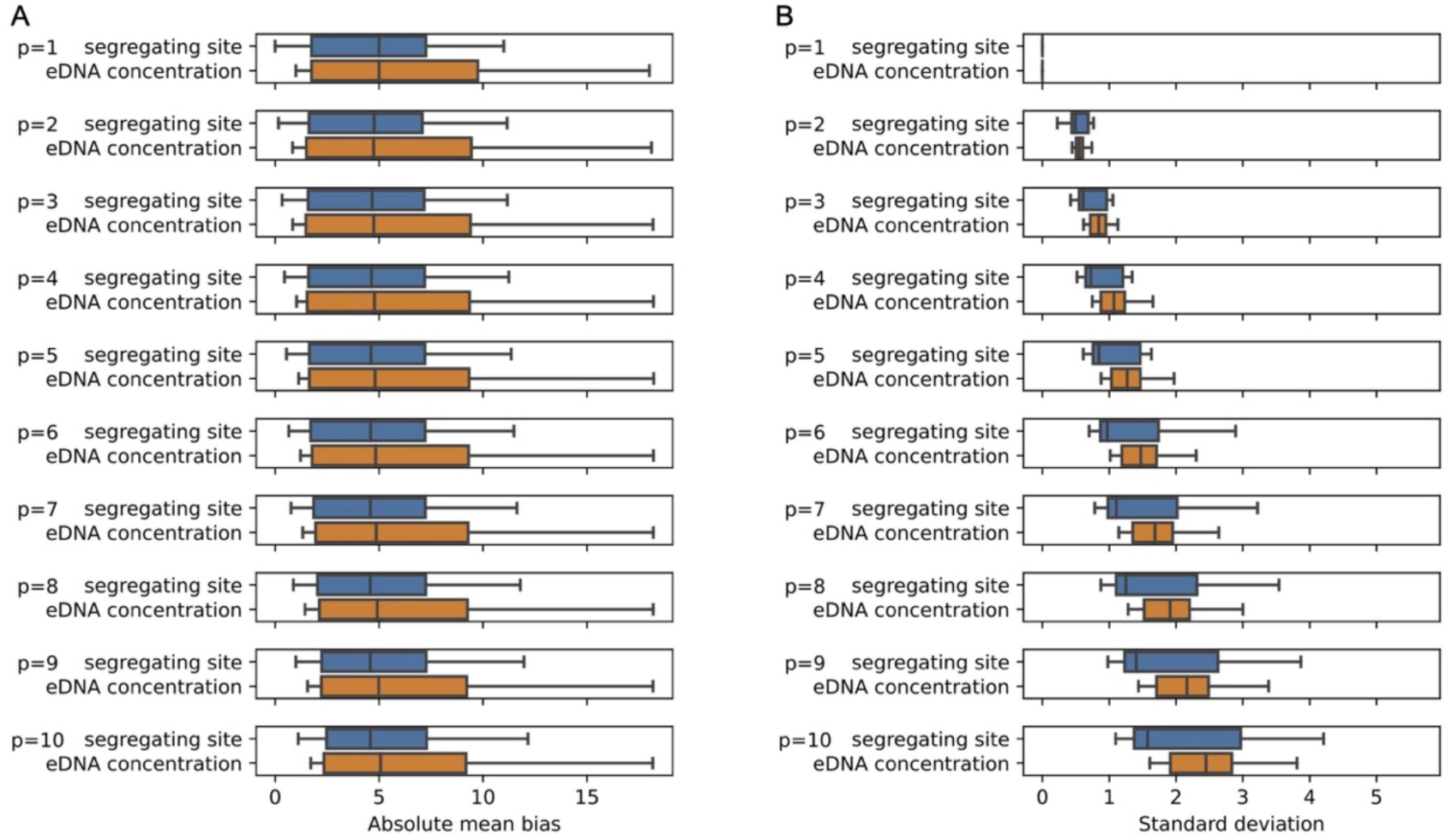
Cross-validation test results for predicting species abundance. Boxplots show (A) absolute mean bias and (B) standard deviation of predicted species abundance based on segregating sites (blue) and eDNA concentration (orange) with increasing size of testing sets (top to bottom, 1 to 10 samples). Lower bias and standard deviation indicate better model accuracy and precision.

## 4 DISCUSSION

Estimating the species abundance in natural ecosystems could provide vital insights into a wide array of evolutionary and ecological questions, and also hold implications for effective conservation strategies. Emerging eDNA technologies, as a non-invasive and rapid approach, represent a promising complement to traditional methods for estimating species abundance. This study proposes using the number of segregating sites in eDNA as a proxy for estimating species abundance. Segregating sites represent inherent genetic variability within a population, making this metric less susceptible to differential individual DNA contributions in eDNA samples than concentration-based methods. Thus, the segregating sites-based approach ensures a more precise and less biased estimate of species abundance. Notably, our study has yet to estimate species abundance in field tests, so the results from the in silico, in vitro, and mesocosm experiments must be interpreted with caution. But still, our study is an essential step toward accurate and precise estimates of species abundance using eDNA.

Previous studies have already leveraged genetic variability as an indicator of species abundance in eDNA samples. Recent methods count distinct microsatellite alleles (Carreon-Martinez et al. 2014) or mtDNA haplotypes (Yoshitake et al. 2019, 2021) as a representation of species abundance. Another approach, the DNA mixture model, uses a probabilistic framework to estimate the number of contributors to eDNA samples based on observed microsatellite alleles or mtDNA haplotypes (Andres et al. 2021, 2023; Sethi et al. 2019). Both methods utilize distinct alleles or haplotypes to infer species abundance but differ in complexity. Allele counting provides a simple, conservative estimate of minimal possible abundance. The DNA mixture model takes this concept further by considering the relative frequency of distinct alleles/haplotypes and accounting for potentially unobserved alleles. In contrast, our method employs a direct correlation between the number of segregating sites, and abundance. This approach simplifies the analytical process, especially compared to the DNA mixture model, while still capturing the underlying relationship between genetic variability and species abundance. Another advantage is that our method does not require sequence assembly when using haplotypes, allowing us to capture variation even when small pieces of sequencing data are missing. To apply our method in the field, a pilot study is required to establish the relationship between the number of segregating sites and species abundance, while theoretically, the allele counting method or DNA mixture model does not need such a pilot study. On the other hand, this initial investigation gives us a good sense of when genetic variability is exhausted, thus helping us avoid scenarios where results might be underestimated without our knowledge.

In the in silico study, we found a strong correlation between genetic variation in sequences and abundance estimates across all tested mutation rates. Notably, data generated under higher mutation rates exhibited stronger correlations between segregating sites and individual counts, with R² values of 0.99, 0.98, and 0.91 for mutation rates of 10⁻⁶/bp/gen, 10⁻⁷/bp/gen, and 10⁻⁸/bp/gen. We also notice that the relationship between segregating sites and species abundance can reach a plateau (Fig. 2), indicating that when genetic variability is exhausted, the maximum detectable species abundance significantly depends on the population’s mutation rate. Our in silico study suggests that even at the lowest mutation rate (10⁻⁸/bp/gen), we can reliably estimate populations of up to 100 individuals, which suffices for most applications (Fig. 2c). However, preliminary surveys are advisable for investigating larger populations.

Complementing the in silico study, the in vitro experiments also demonstrated a strong relationship between the number of segregating sites and species abundance, particularly evident in *A. hasta* (R² = 0.98). However, differing from in silico analysis, neither *A. hasta* nor *T. bifasciatus* showed signs of reaching a plateau in the relationship between segregating sites and abundance. This finding suggests a broader potential for using segregating sites to estimate species abundance in large populations. We noticed lower sequence depth and coverage for enriched data from *T. bifasciatus* in some samples, likely due to excessive PCR cycles during DNA library preparation. This resulted in fewer on-target sequences for *T. bifasciatus* compared to *A. hasta* (Table S2). Despite the relatively poor quality of the *T. bifasciatus* enrichment data, we still identified a strong positive linear relationship between the number of segregating sites and species abundance. This suggests that low sequencing depth did not significantly impact our ability to extract segregating sites.

Mesocosm experiment results showed that the number of individuals predicted the number of segregating sites (R^2^ = 0.70) better than predicting the eDNA concentration (R^2^ = 0.46) as expected. We observed that the overall R² was lower in the mesocosm experiments compared to the in vitro and in silico studies. This difference can be attributed to variables introduced from eDNA samples, such as DNA degradation and the influence of ambient water chemistry on target enrichment. Counter-intuitively, both the number of segregating sites and the number of DNA copies were not heavily affected by body size or feeding/non-feeding treatment in this study, which was inconsistent with previous results (Klymus et al. 2015; Mizumoto et al. 2018). Moreover, eDNA concentrations measured in our study did not show overdispersed distribution as described in the previous study (Chambert et al. 2018). One possible explanation is that the experiment duration was too short, and the behavior of the fish was disrupted, so the size of the fish and their feeding behavior had erratic effects on releasing DNA. Alternatively, unforeseen factors, such as placing aquaria in proximity to corridors or disturbances, may have led to varying stress responses and DNA release from the fish.

Our cross-validation analysis revealed that species abundance estimates using segregating sites outperformed those based on eDNA concentration, showing lower absolute bias and reduced standard deviation. The performance difference became more evident as the testing set size increased. This is because multiple estimates per sample, derived from models built on different training sample sets, provided a more comprehensive representation of the estimate bias and variation distribution. While the segregating sites method showed substantially lower upper quantile absolute mean bias, the difference in median absolute mean bias was less pronounced but still favorable for segregating site methods. We anticipate that longer experiments, which might introduce more confounding factors for the eDNA concentration-based method, could further distinguish the performance of these two approaches.

Nevertheless, a couple of limitations need to be addressed when applying the segregating-site method. Firstly, there are no global criteria for thresholding read depth and minor allele frequency to distinguish polymorphism from sequencing errors. We set an ad hoc criterion of more than seven reads and at least 3% minor-allele reads to call a segregating site. Alternatively, duplex Sequencing (Schmitt et al. 2012) or adding unique molecular identifiers (UMIs) (Yoshitake et al. 2021) may be applied to discriminate genuine polymorphic sites from sequencing errors. Secondly, a pilot survey of the genetic diversity of the population has to be done before extrapolating the correlation between the number of segregating sites and the number of individuals in actual samples. There is often a limited number of individuals sampled in the pilot study. Therefore, future research should explore the optimal number of individuals for the pilot study and how much we can extrapolate the correlation between the number of individuals and segregating sites from those pilot study samples. A recommended approach for estimating the number of individuals needed for a pilot study is to conduct an in silico analysis similar to the one in our study. Researchers can gather relevant parameters for the population of target species, such as mutation rates, from existing literature or by estimating them based on closely related species in the phylogeny. By simulating various scenarios, researchers can obtain a reliable estimate of the sample size required for an effective pilot study.

In this study, we employed target enrichment instead of the conventional metabarcoding approach to collect sequences from eDNA samples. Target enrichment has already been applied in species detection, proving effective in capturing mitochondrial DNA (Jensen et al. 2021; Li et al. 2023; Wilcox et al. 2018) and even nuclear DNA (Jensen et al. 2021) from water samples. Our application extends the utility of target enrichment to species abundance estimates. Compared to metabarcoding, target enrichment is particularly useful when eDNA is partially degraded or when primer binding sites are inaccessible or partially accessible. This advantage is crucial, as non-amplifiable sequences in eDNA could still contain novel alleles or haplotypes not observed in amplifiable sequences, potentially leading to a loss of variability that could bias final species abundance estimates. Moreover, unlike metabarcoding, target enrichment can extract sequences from flanking genomic regions (Li et al. 2013), not just short amplifiable regions suitable for PCR. In our study, this advantage enabled us to obtain segregating sites beyond baits-targeted regions. While previous studies typically designed species-specific probes (Jensen et al. 2021; Li et al. 2023; Wilcox et al. 2018), we designed RNA baits based on conserved mitochondrial regions across vertebrates, allowing our probes to simultaneously capture eDNA from multiple species, thereby reducing costs associated with species-specific baits design.

There are several challenges when applying target enrichment in the field study. eDNA samples typically contain genetic material from a diverse range of species, resulting in the capture of numerous non-target sequences when using the target enrichment method. While mapping to the reference genome can filter out most non-target reads, this approach may still fail to exclude some similar sequences from closely related species that differ by only a few base pairs. This limitation could potentially lead to false positive segregating sites, resulting in inflated species abundance estimates. Another challenge was the variability in capture efficiency. Although most eDNA samples achieved satisfactory sequencing depth (over 50x) (Table S3), some samples exhibited poor capture efficiency. For instance, one sample, FW1N5, only attained an average sequencing depth of 18x, which might impede the detection of rare segregating sites. These limitations underscore the need for careful consideration and potential refinement of target enrichment protocols when applied to eDNA samples.

eDNA techniques provide a non-invasive and effective solution for biomonitoring, providing insight into ecological and evolutionary processes in the wild. Qualitative studies, such as species detection, are well-established using eDNA, while quantitative studies that require accurate species abundance estimates still pose challenges. Our study proposes the use of segregating site counts as a more precise and less biased proxy than eDNA concentration, marking a significant advance toward accurate species abundance estimates from eDNA. The following steps in developing this method will include (1) accurately identifying rare alleles with low frequencies from eDNA data, (2) extended testing of different species in mesocosms and the fields for longer durations, and (3) investigating effective sampling strategies for pilot studies.

## AUTHOR CONTRIBUTIONS

Q.A. and C.L. designed the study. Q.A., H.Y., and Y.W. conducted the study and collected specimens. Q.A. and H.Y. completed laboratory work, analyzed the data, and interpreted the results. Q.A. and H.Y. wrote the manuscript. C.L. provided feedback on the manuscript. All authors approved the manuscript for publication.

## ACKNOWLEDGEMENTS

This work was supported by the National Key Research and Development Program of China (2022YFC2601301). Dr Z. Zhang helped with model selection and correlation analyses.

## DATA ACCESSIBILITY STATEMENT

Simulated sequences for in-silico study are available at https://doi.org/10.5061/dryad.w3r2280zz.

Raw sequences of targeted enrichment from tissues of *A. hasta* and *T. bifasciatus* were deposited in NCBI’s Sequence Read Archive (BioProject ID: PRJNA1100443). Raw sequences of enriched sequences from eDNA samples were also available at NCBI’s Sequence Read Archive (BioProject ID: PRJNA1100453).

## BENEFIT-SHARING STATEMENT

The study developed an accurate and precise method for estimating species abundance using eDNA data. This advancement will guide future efforts in non-invasive biomonitoring techniques for conservation ecology and population genetics. All data from the study are freely accessible through the NCBI’s Sequence Read Archive and DRYAD digital repository.

---

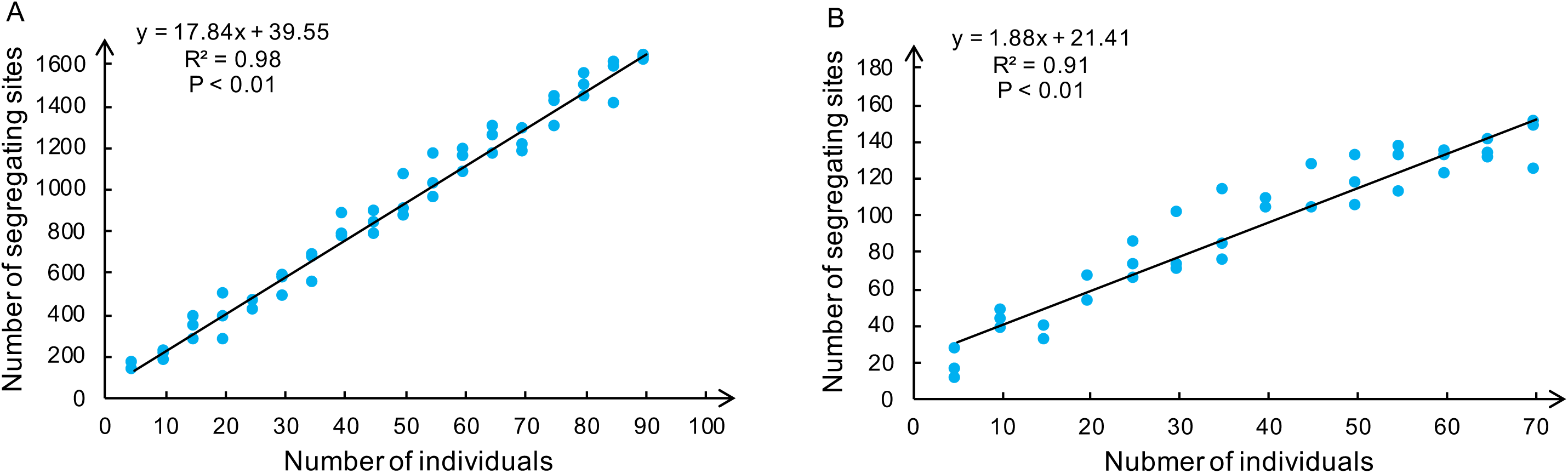

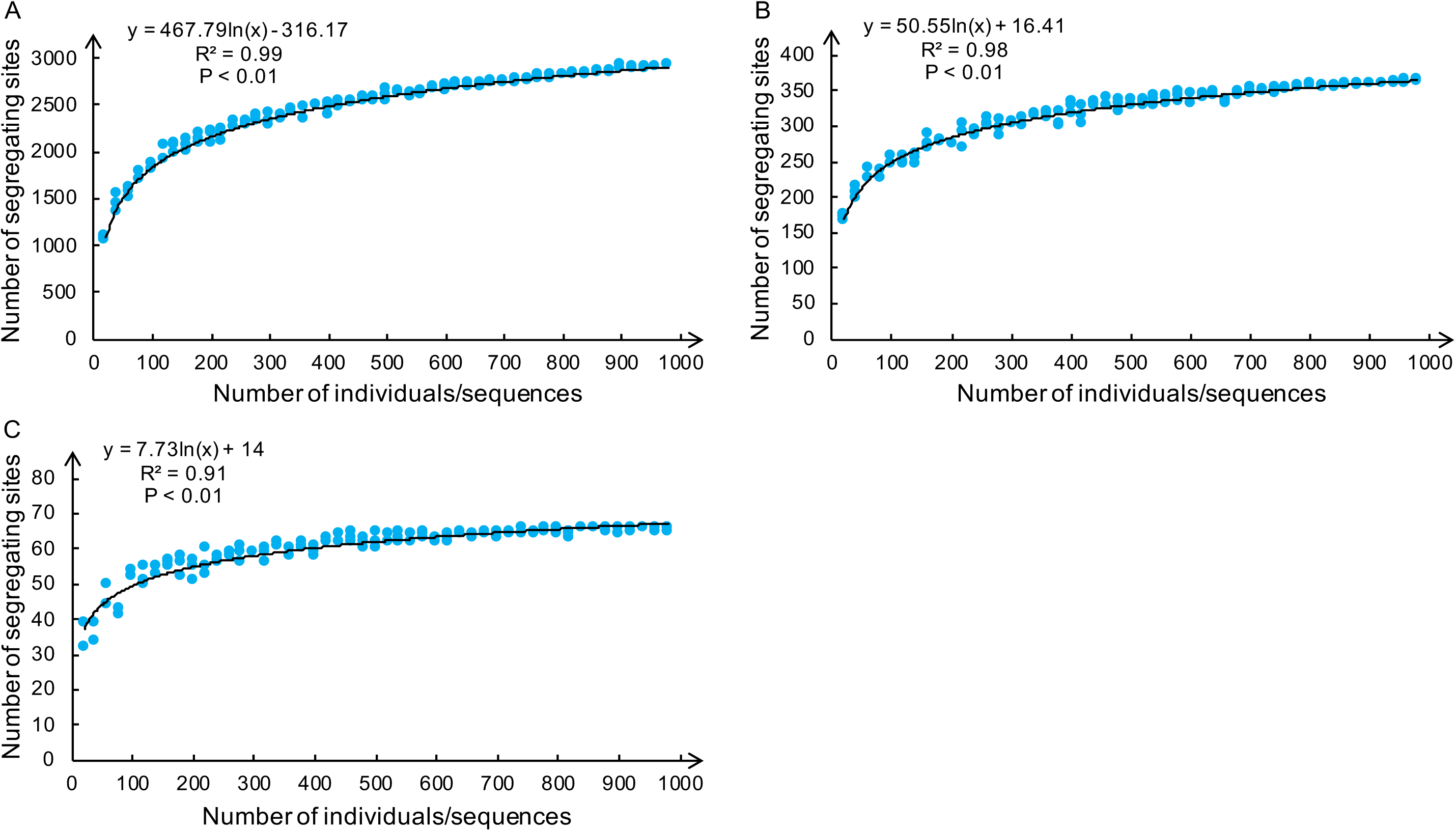

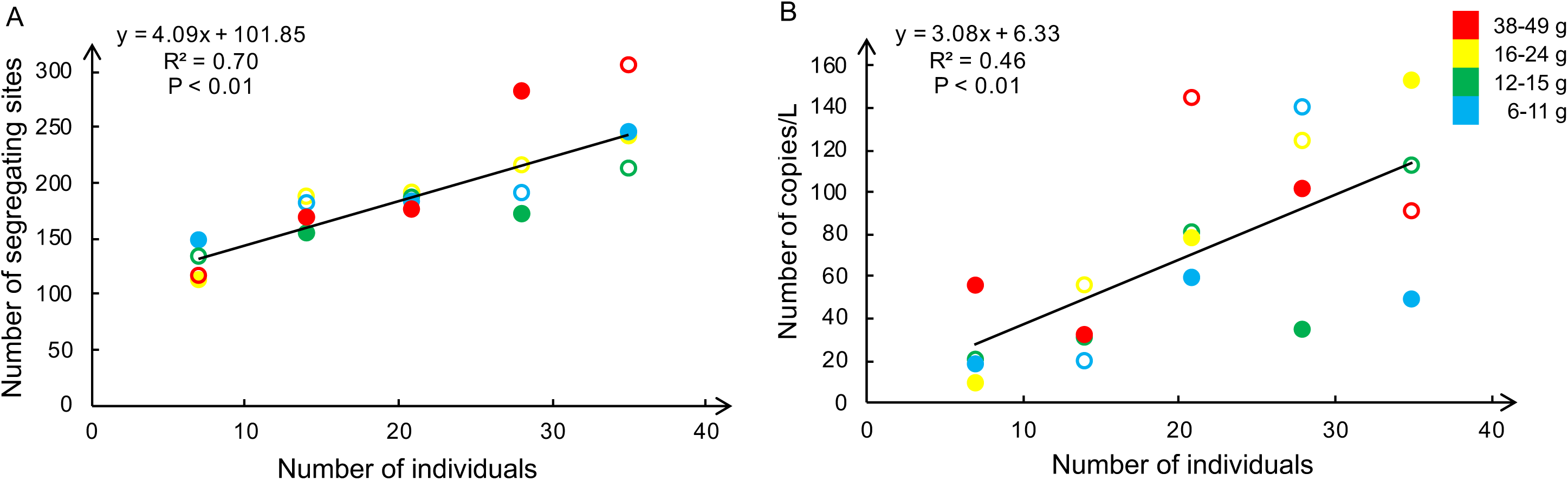

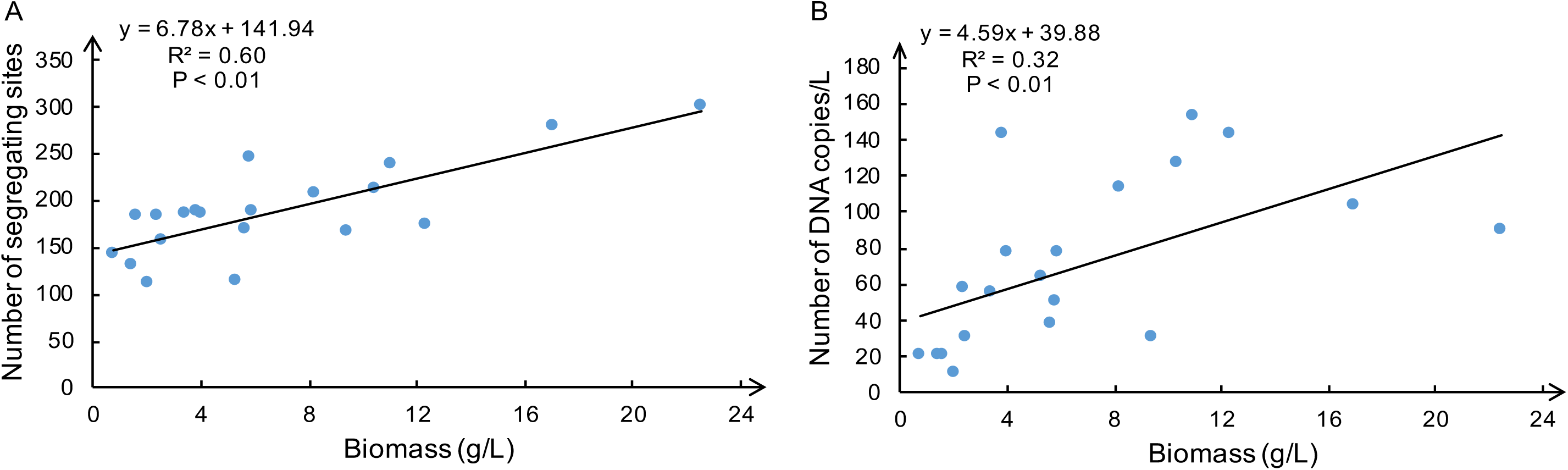

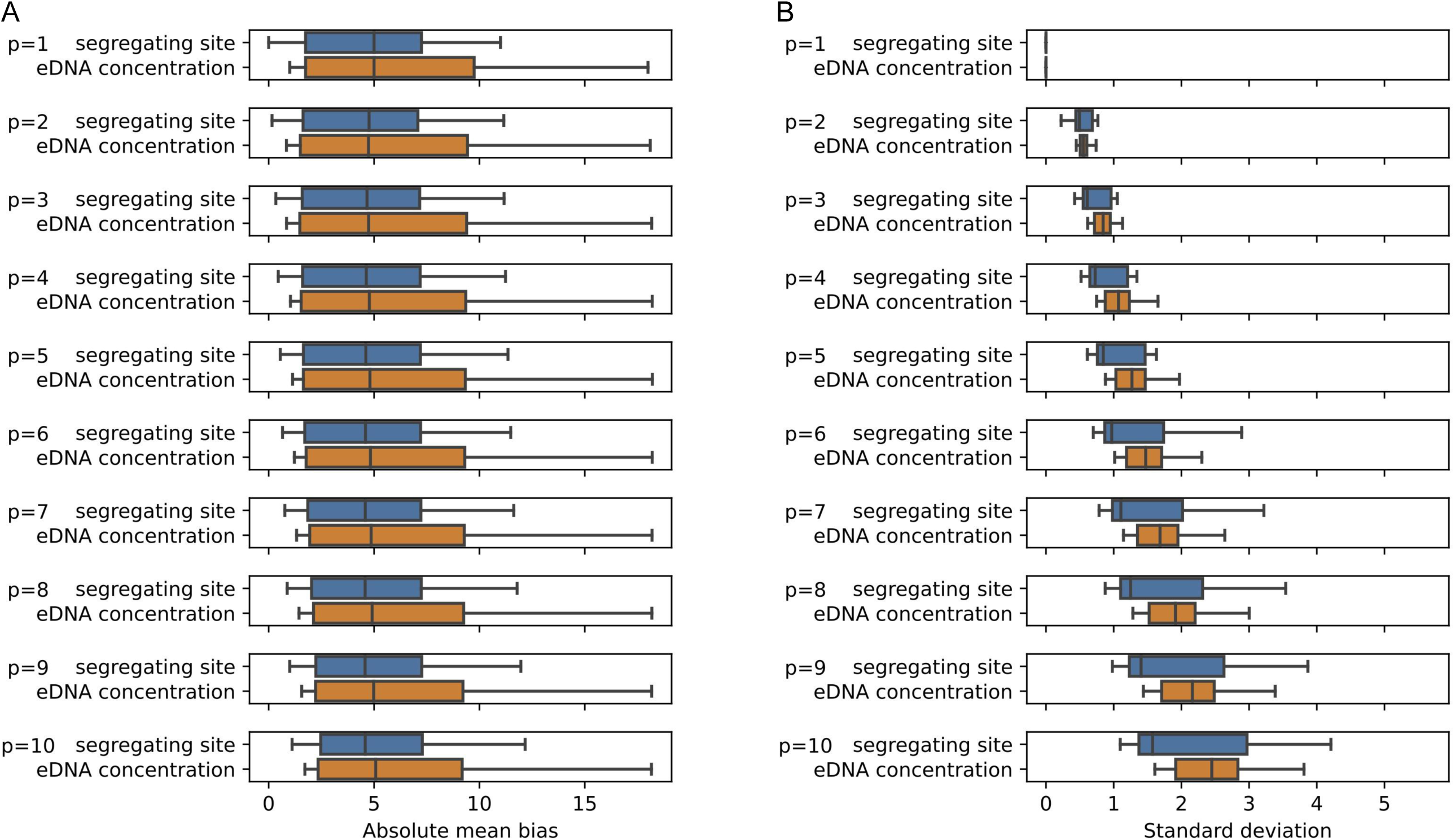

