## Supplementary file for "Estimation of Species Abundance Based on the Number of Segregating Sites using Environmental DNA (eDNA)"

**Species Abundance can be Better Estimated from the Number of Segregating Sites using Environmental DNA (eDNA)**

**Supplementary materials:**

Fig. S1

Fig. S2

Table S1

Table S2

Table S3

Table S4

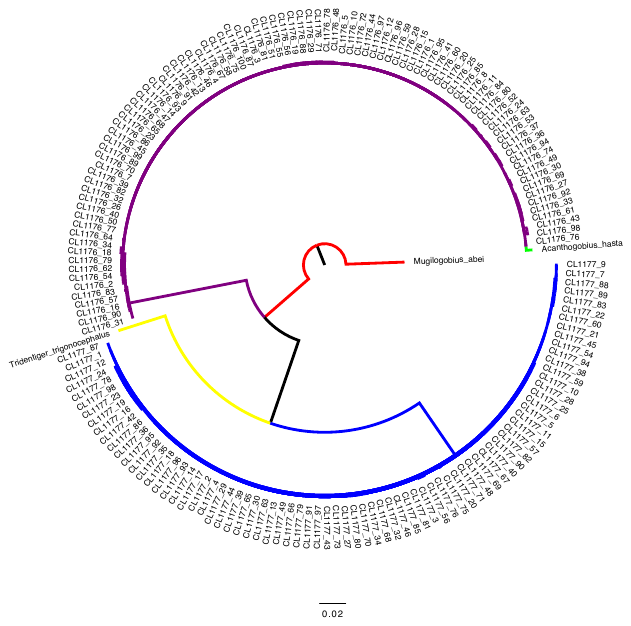

**Fig. S1.** Maximum likelihood (RAxML) tree of *Acanthogobius hasta* (purple) and *Tridentiger bifasciatus* (blue) based on 11 loci. *Mugilogobious abei* (red) was used as the outgroup. Sequences of *Tridentiger trigonocephalus* (yellow) and *Acanthogobius hasta* (green) retrieved from the Genbank were used for comparison.

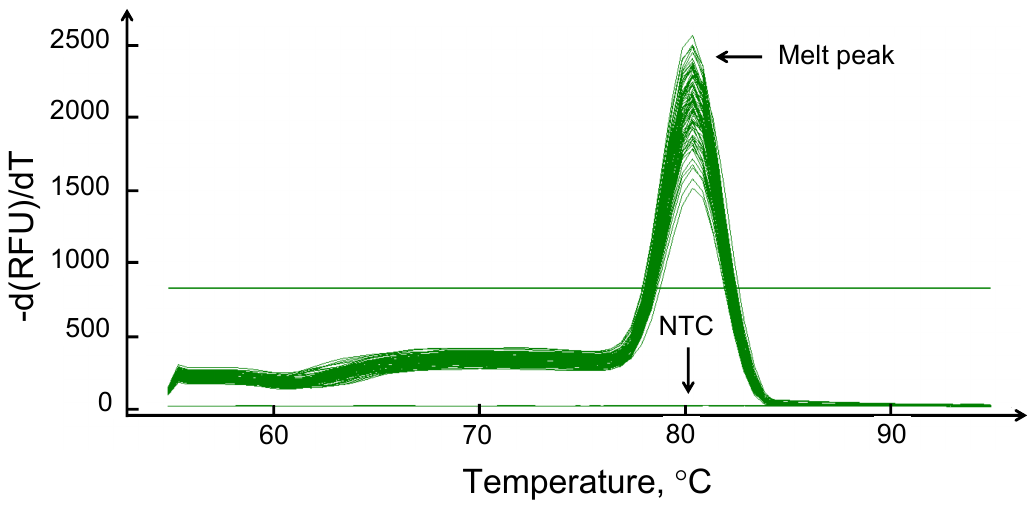

**Fig. S2.** Melt curves of qPCR amplifying the T2 target. Each melt curve represented one sample. All curves had a single peak at 80 °C except the three no-template control (NTC) samples.

**Table S1.** Mitochondrial sequences of *Acanthogobius hasta* used for RNA baits design

| Targets | Sequences |
| --- | --- |
| T1 | TTAAAACCCAAAGGACCTGGCGGTGCTTTAGACCCCCCTAGAGGAGCCTGTTCTATAACCGATAATCCCCGTTAAACCTCACCCTCCCTTGTTCTCCCCGCCTATATACCGCCGTCGTCA |
| T2 | GCCGCGGTATCCTGACCGTGCAAAGGTAGCGTAATCACTTGTCTTTTAAATGGAGACCTGTATGAATGGCAAGACGAGGGCTAAACTGTCTCCTTTTTCCAGTCAATGAAATTGATCTCC |
| T3 | GTTTACGACCTCGATGTTGGATCAGGACATCCTAATGGTGCAGCCGCTATTAAGGGTTCGTTTGTTCAACGATTAAAGTCCTACGTGATCTGAGTTCAGACCGGAGAAATCCAGGTCAGT |
| T4 | ACCCTACCTGAAGAGATCAAAACTCTTAGTGCTTCCACTACACCACTTCCTAGTAAAGTCCGCTAAATAAGCTTTTGGGCCCATACCCCAAACATGATGGTTAAACCCCCTCCTTTGCT |
| T5 | CTTCTGCGTGCAAAGCAGACACTTTAATTAAGCTACAACCTTTTTCTAGACAGGAAGGCCTCGATCCTACAAACTCTTAGTTAACAGCTAAGCGCCCAAACCAGGCGGGCATCCGCCTACTCT |
| T6 | TGGCAATTACACGTTGATTTTTCTCGACAAACCATAAAGACATTGGCACCCTCTATCTTGTATTTGGTGCCTGAGCCGGAATAGTAGGCACAGCTCTAAGCCTTTTAATTCGGGCTGAAC |
| T7 | AATAATTTTCTTTATAGTAATACCAATCATAATTGGAGGGTTTGGGAACTGACTAATCCCCCTAATGATTGGTGCCCCTGATATAGCCTTTCCCCGAATAAACAACATAAGCTTTTGACT |
| T8 | CACTATGCTTCTAACAGATCGCAACCTAAACACATCCTTCTTTGACCCTGCAGGGGGTGGTGACCCGATTCTTTACCAACACCTTTTCTGATTTTTTGGCCATCCAGAAGTCTACATTCT |
| T9 | TATTGTTTGAGCACATCATATGTTCACCGTAGGAATGGACGTTGACACACGGGCATACTTCACCTCCGCAACAATAATTATTGCCATCCCTACAGGTGTAAAAGTCTTTAGCTGACTAGC |
| T10 | GCAGTTGCCATGATTCAAGCATACGTCTTCATTCTTCTAATTAGCCTCTATCTTCAAGAAAACGTCTAATGGCCCGCCAAGCACACGCATTTCACATAGTAGACCTTAGCCCCTGACCTCC |
| T11 | GACCCAGACAGTTTTACCCCTGCCAACCCGCTTGTTACTCCACCACACATTAAGCCAGAGTGATACTTCTTATTTGCCTACGCCATCTTACGCTCCATTCCGAATAAACTCGGCGGAGT |

**Table S2.** Sequencing results for the individual fish samples

| Voucher number | Raw reads  (bp) | Trimmed reads  (bp) | De-duplicated reads  (bp) | On-target reads (%) | Seq. depth |
| --- | --- | --- | --- | --- | --- |
| CL1176_1 | 25,311,812 | 24,486,338 | 6,994,878 | 29,181 (11.75) | 208 |
| CL1176_2 | 22,124,454 | 21,454,444 | 5,020,882 | 16,536 (7.61) | 122 |
| CL1176_3 | 10,036,168 | 9,729,973 | 2,633,984 | 8,757 (8.88) | 67 |
| CL1176_4 | 10,639,946 | 10,290,246 | 3,292,388 | 11,461 (10.98) | 83 |
| CL1176_5 | 11,130,604 | 10,692,057 | 2,819,979 | 5,425 (4.96) | 44 |
| CL1176_7 | 15,177,876 | 14,644,671 | 3,941,647 | 15,542 (10.44) | 112 |
| CL1176_8 | 20,878,114 | 20,158,977 | 5,883,941 | 18,247 (8.91) | 130 |
| CL1176_9 | 27,131,428 | 26,187,774 | 6,352,799 | 18,396 (6.93) | 139 |
| CL1176_10 | 10,770,236 | 10,371,836 | 2,751,483 | 7,226 (6.84) | 55 |
| CL1176_11 | 8,830,834 | 8,550,058 | 2,521,070 | 6,551 (7.55) | 50 |
| CL1176_12 | 3,517,628 | 3,332,711 | 1,009,333 | 1,616 (4.7) | 18 |
| CL1176_13 | 9,627,926 | 9,304,935 | 2,860,585 | 7,475 (7.91) | 57 |
| CL1176_14 | 10,398,354 | 10,024,128 | 3,400,875 | 8,683 (8.52) | 67 |
| CL1176_15 | 12,921,738 | 12,421,125 | 3,758,888 | 1,1542 (9.13) | 85 |
| CL1176_16 | 13,052,230 | 12,553,103 | 3,642,945 | 8,279 (6.47) | 62 |
| CL1176_18 | 11,220,898 | 10,864,256 | 3,440,493 | 11,049 (10.02) | 80 |
| CL1176_19 | 14,816,902 | 14,356,104 | 4,575,403 | 18,029 (12.39) | 132 |
| CL1176_20 | 9,167,164 | 8,883,500 | 2,502,900 | 6,670 (7.4) | 53 |
| CL1176_23 | 4,337,748 | 4,145,146 | 1,424,123 | 1,905 (4.46) | 18 |
| CL1176_24 | 10,312,504 | 9,996,035 | 3,126,951 | 7,884 (7.78) | 61 |
| CL1176_25 | 8,202,210 | 7,953,944 | 1,979,085 | 6,773 (8.39) | 55 |
| CL1176_26 | N/A | N/A | N/A | N/A | N/A |
| CL1176_27 | N/A | N/A | N/A | N/A | N/A |
| CL1176_28 | N/A | N/A | N/A | N/A | N/A |
| CL1176_29 | N/A | N/A | N/A | N/A | N/A |
| CL1176_30 | 10,146,864 | 9,841,738 | 2,504,733 | 9,452 (9.47) | 69 |
| CL1176_31 | 1,883,852 | 1,787,273 | 576,640 | 780 (4.22) | 9 |
| CL1176_32 | 12,013,546 | 11,603,285 | 3,333,553 | 8,727 (7.39) | 67 |
| CL1176_33 | 18,362,810 | 17,790,479 | 5,225,842 | 15,093 (8.36) | 113 |
| CL1176_34 | 19,392,000 | 18,805,176 | 4,655,649 | 16,178 (8.48) | 119 |
| CL1176_36 | 8,610,250 | 8,343,541 | 2,502,138 | 7,667 (9.06) | 57 |
| CL1176_37 | 4,037,778 | 3,919,903 | 1,158,951 | 3,748 (9.45) | 31 |
| CL1176_39 | 7,136,458 | 6,923,464 | 1,985,711 | 7,715 (11) | 59 |
| CL1176_40 | 6,184,634 | 5,987,380 | 1,464,959 | 6,001 (9.87) | 48 |
| CL1176_41 | 6,200,188 | 6,004,258 | 1,932,543 | 6,638 (10.9) | 50 |
| CL1176_42 | 59,896,636 | 57,870,375 | 17,103,660 | 51,929 (8.84) | 370 |
| CL1176_43 | 6,002,430 | 5,789,911 | 1,771,426 | 5,309 (9.01) | 40 |
| CL1176_44 | 13,081,722 | 12,603,688 | 4,278,126 | 9,822 (7.65) | 74 |
| CL1176_45 | 10,023,240 | 9,588,100 | 4,108,573 | 6,763 (6.89) | 51 |
| CL1176_46 | 5,421,478 | 5,060,250 | 1,802,657 | 8,167 (15.63) | 62 |
| CL1176_47 | 5,861,838 | 5,667,766 | 1,888,703 | 4,984 (8.67) | 39 |
| CL1176_48 | 4,413,094 | 4,226,831 | 1,620,874 | 3,508 (8.12) | 29 |
| CL1176_49 | 3,556,614 | 3,383,664 | 1,090,666 | 1,932 (5.54) | 17 |
| CL1176_50 | 13,166,764 | 12,778,452 | 3,582,072 | 8,920 (6.88) | 68 |
| CL1176_51 | 32,915,496 | 31,913,161 | 9,042,327 | 25,953 (8.02) | 185 |
| CL1176_52 | 29,017,502 | 28,171,995 | 8,812,769 | 27,924 (9.78) | 199 |
| CL1176_53 | 21,799,436 | 21,191,193 | 6,164,401 | 18,866 (8.79) | 138 |
| CL1176_54 | 22,047,896 | 21,306,586 | 6,594,275 | 20,876 (9.65) | 151 |
| CL1176_55 | 23,623,900 | 22,864,809 | 7,506,455 | 20,818 (8.97) | 147 |
| CL1176_56 | 16,241,002 | 15,721,149 | 5,105,233 | 15,375 (9.64) | 110 |
| CL1176_57 | 23,583,904 | 22,815,232 | 6,787,143 | 18,849 (8.15) | 135 |
| CL1176_58 | 22,782,368 | 22,042,911 | 6,943,261 | 22,842 (10.2) | 165 |
| CL1176_59 | 12,969,410 | 12,550,403 | 4,828,474 | 9,818 (7.71) | 73 |
| CL1176_60 | N/A | N/A | N/A | N/A | N/A |
| CL1176_61 | N/A | N/A | N/A | N/A | N/A |
| CL1176_62 | N/A | N/A | N/A | N/A | N/A |
| CL1176_63 | 21,338,270 | 20,652,082 | 5,758,953 | 17,315 (8.27) | 126 |
| CL1176_64 | 38,592,100 | 37,409,192 | 10,349,381 | 29,573 (7.80) | 211 |
| CL1176_65 | 40,018,018 | 38,773,889 | 9,809,786 | 38,814 (9.87) | 276 |
| CL1176_67 | 33,657,846 | 32,552,951 | 10,770,508 | 30,887 (9.35) | 218 |
| CL1176_68 | 25,695,612 | 24,682,325 | 8,355,044 | 14,255 (5.67) | 109 |
| CL1176_69 | 20,913,464 | 20,147,259 | 8,838,246 | 16,622 (8.11) | 120 |
| CL1176_70 | 15,890,330 | 15,203,693 | 7,162,953 | 10,851 (7.01) | 80 |
| CL1176_71 | 33,874,592 | 32,783,659 | 13,070,729 | 20,621 (6.2) | 152 |
| CL1176_72 | 22,818,324 | 22,092,123 | 6,021,729 | 24,262 (10.84) | 175 |
| CL1176_74 | 35,217,084 | 34,207,475 | 7,581,164 | 28,726 (8.29) | 207 |
| CL1176_75 | 22,652,886 | 21,982,076 | 6,633,642 | 18,063 (8.11) | 131 |
| CL1176_76 | 30,523,210 | 29,611,376 | 8,669,262 | 23,381 (7.79) | 169 |
| CL1176_77 | 16,829,024 | 16,349,287 | 4,967,676 | 15,995 (9.66) | 119 |
| CL1176_78 | 25,961,848 | 25,256,276 | 6,679,100 | 23,835 (9.33) | 172 |
| CL1176_79 | 21,548,552 | 20,950,075 | 6,240,058 | 16,779 (7.91) | 124 |
| CL1176_80 | 22,396,144 | 21,761,669 | 6,346,111 | 20,270 (9.21) | 148 |
| CL1176_81 | 15,457,242 | 15,005,138 | 3,503,776 | 15,508 (10.21) | 113 |
| CL1176_82 | 21,887,912 | 21,327,737 | 5,581,876 | 16,442 (7.62) | 123 |
| CL1176_83 | 13,635,808 | 13,239,278 | 3,972,252 | 12,521 (9.34) | 91 |
| CL1176_84 | 19,289,990 | 18,705,648 | 5,552,055 | 15,843 (8.35) | 116 |
| CL1176_85 | 18,620,158 | 18,044,978 | 5,967,945 | 18,977 (10.38) | 137 |
| CL1176_86 | 26,396,350 | 25,387,107 | 8,160,149 | 23,042 (8.91) | 165 |
| CL1176_87 | 19,977,598 | 19,413,728 | 5,358,639 | 16,433 (8.36) | 121 |
| CL1176_88 | 17,827,106 | 17,214,881 | 5,197,089 | 17,429 (9.98) | 125 |
| CL1176_89 | 20,493,304 | 19,848,482 | 5,986,870 | 21,721 (10.79) | 157 |
| CL1176_90 | 18,486,232 | 17,932,734 | 5,389,006 | 19,645 (10.81) | 140 |
| CL1176_91 | 45,405,762 | 44,213,074 | 9,451,015 | 48,345 (10.81) | 344 |
| CL1176_92 | 18,378,364 | 17,803,724 | 5,638,073 | 18,322 (10.14) | 130 |
| CL1176_93 | 25,253,434 | 24,454,510 | 7,398,151 | 23,804 (9.59) | 169 |
| CL1176_94 | 28,840,348 | 27,947,732 | 7,820,975 | 25,113 (8.86) | 181 |
| CL1176_95 | 29,461,700 | 28,552,310 | 8,617,998 | 24,380 (8.42) | 175 |
| CL1176_96 | 32,485,640 | 31,609,801 | 7,806,435 | 28,897 (9.03) | 210 |
| CL1176_97 | 16,472,292 | 15,852,722 | 5,015,248 | 22,392 (13.88) | 160 |
| CL1176_98 | 16,539,356 | 15,974,942 | 4,362,344 | 18,972 (11.7) | 135 |
| CL1176_99 | 12,543,998 | 12,137,014 | 4,799,371 | 16,491 (13.39) | 119 |
| CL1176_100 | 35,865,504 | 34,729,435 | 9,768,487 | 43,505 (12.35) | 308 |
| Average | 18,485,003 | 17,891,350 | 5,304,806 | 16,354 (9) | 119 |
| CL1177_1 | 103,285,832 | 98,359,828 | 207,193 | 25,636 (2.54) | 178 |
| CL1177_2 | 33,004,578 | 31,246,344 | 56,140 | 5,451 (1.69) | 40 |
| CL1177_3 | 26,528,054 | 25,251,784 | 46,653 | 4,355 (1.69) | 33 |
| CL1177_4 | 11,099,294 | 10,554,312 | 17,386 | 1,636 (1.5) | 13 |
| CL1177_5 | 56,857,344 | 54,343,432 | 96,794 | 9,816 (1.77) | 68 |
| CL1177_6 | 70,332,360 | 67,223,065 | 101,870 | 10,980 (1.6) | 77 |
| CL1177_7 | 50,746,440 | 48,296,656 | 98,276 | 10,472 (2.12) | 73 |
| CL1177_9 | 41,251,632 | 38,837,604 | 166,614 | 16,356 (1.81) | 114 |
| CL1177_10 | 91,589,022 | 87,747,992 | 166,143 | 17,217 (1.92) | 119 |
| CL1177_11 | 89,982,920 | 86,142,379 | 118,199 | 11,809 (1.34) | 82 |
| CL1177_12 | 36,892,068 | 35,153,355 | 57,429 | 6,856 (1.9) | 48 |
| CL1177_13 | 62,123,686 | 59,416,480 | 113,295 | 11,007 (1.81) | 77 |
| CL1177_14 | 47,525,752 | 44,949,913 | 71,792 | 8,539 (1.85) | 59 |
| CL1177_15 | 62,714,738 | 60,024,755 | 105,777 | 10,574 (1.72) | 73 |
| CL1177_16 | 69,296,908 | 66,698,483 | 146,235 | 13,447 (1.98) | 93 |
| CL1177_17 | 42,683,206 | 40,888,733 | 77,870 | 8,429 (2.02) | 60 |
| CL1177_18 | 42,449,088 | 40,577,739 | 47,612 | 5,830 (1.4) | 41 |
| CL1177_19 | 47,459,092 | 45,434,630 | 67,124 | 7,039 (1.52) | 50 |
| CL1177_20 | 86,039,274 | 82,046,298 | 154,350 | 15,271 (1.82) | 106 |
| CL1177_21 | 471,829,58 | 449,185,24 | 78,932 | 9,171 (1.99) | 64 |
| CL1177_22 | 38,803,392 | 36,787,179 | 73,649 | 8,401 (2.22) | 58 |
| CL1177_23 | 63,501,326 | 60,607,866 | 73,489 | 6,909 (1.11) | 49 |
| CL1177_24 | 47,018,732 | 44,859,771 | 99,377 | 10,531 (2.29) | 73 |
| CL1177_25 | 53,172,864 | 50,630,171 | 106,048 | 11,427 (2.2) | 79 |
| CL1177_27 | 9,899,818 | 9,152,133 | 13,257 | 1,396 (1.45) | 12 |
| CL1177_28 | 30,257,176 | 29,078,130 | 58,463 | 6,376 (2.14) | 46 |
| CL1177_29 | 16,534,508 | 15,477,946 | 17,211 | 1,561 (0.97) | 14 |
| CL1177_30 | 499,748 | 453,072 | 291 | 45 (0.94) | 1 |
| CL1177_32 | 799,112 | 727,739 | 1,079 | 64 (0.83) | 2 |
| CL1177_34 | 407,838 | 372,504 | 522 | 66 (1.67) | 2 |
| CL1177_35 | 26,337,568 | 24,640,851 | 45,028 | 4,796 (1.89) | 35 |
| CL1177_36 | 7,210,794 | 6,470,257 | 7,770 | 705 (1.02) | 7 |
| CL1177_38 | 21,456,440 | 20,409,847 | 41,875 | 3,860 (1.84) | 29 |
| CL1177_39 | 332,492 | 290,450 | 370 | 50 (1.59) | 2 |
| CL1177_40 | 11,223,322 | 10,658,835 | 15,540 | 1,471 (1.34) | 12 |
| CL1177_42 | 10,161,408 | 9,125,250 | 10,460 | 757 (0.78) | 7 |
| CL1177_43 | 9,665,498 | 9,012,774 | 13,937 | 1,170 (1.24) | 10 |
| CL1177_44 | 16,163,232 | 15,174,093 | 18,204 | 1,436 (0.91) | 12 |
| CL1177_45 | 5,871,534 | 5,384,695 | 6,065 | 1,311 (2.31) | 11 |
| CL1177_46 | 1,877,186 | 1,715,980 | 1,224 | 151 (0.83) | 3 |
| CL1177_48 | 1,711,546 | 1,565,209 | 1,360 | 145 (0.87) | 2 |
| CL1177_49 | 285,426 | 242,638 | 99 | 42 (1.54) | 2 |
| CL1177_54 | 239,376,262 | 223,470,776 | 13,297 | 1,314 (0.06) | 11 |
| CL1177_56 | N/A | N/A | N/A | N/A | N/A |
| CL1177_57 | N/A | N/A | N/A | N/A | N/A |
| CL1177_59 | 11,294,426 | 10,537,320 | 12,362 | 1,231 (1.12) | 11 |
| CL1177_60 | N/A | N/A | N/A | N/A | N/A |
| CL1177_63 | 637,512 | 545,837 | 406 | 65 (1.06) | 2 |
| CL1177_65 | 248,258 | 222,326 | 142 | 54 (2.24) | 2 |
| CL1177_66 | 3,309,770 | 2,857,828 | 3,788 | 304 (0.99) | 4 |
| CL1177_67 | 8,274,526 | 7,473,098 | 8,321 | 758 (0.96) | 7 |
| CL1177_68 | 800,122 | 701,929 | 644 | 52 (0.68) | 2 |
| CL1177_69 | 7,591,160 | 7,106,402 | 4,566 | 392 (0.53) | 5 |
| CL1177_70 | 3,313,002 | 3,114,949 | 518 | 85 (0.26) | 2 |
| CL1177_71 | 2,473,692 | 2,207,645 | 1,468 | 155 (0.65) | 2 |
| CL1177_73 | 11,002,536 | 10,410,848 | 16,636 | 1,488 (1.38) | 13 |
| CL1177_75 | 10,287,254 | 9,338,252 | 11,325 | 1,023 (1.03) | 9 |
| CL1177_76 | 1,959,198 | 1,729,933 | 2,471 | 200 (1.08) | 3 |
| CL1177_78 | 7,057,072 | 6,597,247 | 11,164 | 947 (1.37) | 9 |
| CL1177_79 | 1,919,000 | 1,752,236 | 2,217 | 192 (1.04) | 3 |
| CL1177_80 | 5,215,034 | 4,884,944 | 5,697 | 561 (1.1) | 6 |
| CL1177_81 | 748,006 | 647,251 | 620 | 62 (0.86) | 2 |
| CL1177_82 | 4,093,934 | 3,844,484 | 4,292 | 441 (1.1) | 5 |
| CL1177_83 | 15,522,488 | 14,590,606 | 13,718 | 1,381 (0.91) | 11 |
| CL1177_85 | 3,084,136 | 2,692,214 | 2,396 | 222 (0.75) | 3 |
| CL1177_86 | 3,899,812 | 3,467,325 | 3,331 | 301 (0.81) | 3 |
| CL1177_87 | 12,676,914 | 12,035,359 | 16,383 | 1,555 (1.25) | 13 |
| CL1177_88 | 35,093,460 | 33,131,524 | 57,448 | 6,634 (1.94) | 46 |
| CL1177_89 | 37,759,860 | 35,686,070 | 56,995 | 6,673 (1.82) | 47 |
| CL1177_90 | 40,578,972 | 38,556,685 | 66,601 | 6,817 (1.72) | 48 |
| CL1177_91 | 28,457,154 | 26,997,883 | 54,664 | 6,240 (2.25) | 44 |
| CL1177_92 | 40,827,230 | 38,574,479 | 57,970 | 6,497 (1.63) | 45 |
| CL1177_93 | 41,750,774 | 39,726,140 | 80,210 | 8,134 (1.99) | 57 |
| CL1177_94 | 41,251,632 | 38,837,604 | 56,455 | 5,709 (1.43) | 41 |
| CL1177_95 | 31,132,644 | 29,550,575 | 44,634 | 4783 (1.58) | 34 |
| CL1177_96 | 34,022,254 | 32,378,502 | 61,774 | 6,883 (2.07) | 48 |
| CL1177_97 | 79,545,782 | 74,652,066 | 83,875 | 8,333 (1.08) | 58 |
| CL1177_98 | 76,062,898 | 71,242,273 | 119,234 | 10,116 (1.39) | 73 |
| Average | 30,896,203 | 29,264,781 | 46,355 | 4,829 (1.44) | 35 |

**Table S3.** Sequencing results for the environment DNA samples

| Sample | Raw Reads | Trimmed Reads | De-duplicated Reads | On-target Reads (%) | Seq. Depth |
| --- | --- | --- | --- | --- | --- |
| F0W3N4 | 941,238,144 | 935,387,271 | 42,187,358 | 55,743 (0.85) | 483 |
| F0W2N3 | 817,438,464 | 812,575,293 | 124,645,415 | 35,027 (0.62) | 332 |
| F0W4N1 | 644,261,184 | 639,486,317 | 161,928,500 | 33,198 (0.74) | 305 |
| FW3N3 | 738,235,584 | 733,108,832 | 138,430,850 | 29,683 (0.58) | 286 |
| F0W2N5 | 709,793,280 | 705,333,341 | 143,540,357 | 26,973 (0.55) | 240 |
| F0W3N2 | 575,535,168 | 571,603,775 | 119,572,288 | 21,858 (0.55) | 197 |
| FW2N2 | 468,889,716 | 465,133,668 | 221,937,506 | 28,501 (0.88) | 301 |
| F0W2N1 | 421,112,028 | 418,007,283 | 206,231,492 | 22,273 (0.77) | 233 |
| FW2N4 | 413,983,212 | 410,750,790 | 182,225,859 | 24,609 (0.86) | 245 |
| FW1N1 | 490,915,092 | 487,310,277 | 178,334,431 | 19,537 (0.58) | 198 |
| F0W4N3 | 464,080,236 | 460,118,951 | 225,600,484 | 22,254 (0.7) | 232 |
| FW4N2 | N/A | N/A | N/A | N/A | N/A |
| FW3N1 | 288,058,824 | 286,112,260 | 150,056,804 | 16,320 (0.82) | 180 |
| F0W1N2 | 286,215,756 | 283,874,318 | 143,891,787 | 14,802 (0.75) | 167 |
| F0W1N4 | 339,080,520 | 334,551,555 | 165,523,789 | 11,525 (0.5) | 99 |
| FW1N3 | 434,562,492 | 430,667,565 | 254,254,537 | 8,857 (0.3) | 96 |
| FW4N4 | 289,893,372 | 286,787,688 | 144,707,516 | 7,548 (0.39) | 50 |
| FW3N5 | 428,823,708 | 423,960,855 | 164,177,350 | 5,928 (0.21) | 53 |
| F0W4N5 | 1,047,420,684 | 1,036,575,890 | 227,387,844 | 4,202 (0.06) | 36 |
| FW1N5 | 589,433,940 | 582,598,631 | 124,126,132 | 2,009 (0.05) | 18 |
| Average | 546,787,969 | 542,312,872 | 164,145,279 | 20,571 (0.59) | 197 |

**Table S4.** Primers for amplifying tissue samples (*Tridentiger bifasciatus* and *Acanthogobius hasta*) and for making home-made gene-capture baits (*Acanthogobius hasta*)

| Primer Name | *Tridentiger bifasciatus* | *Acanthogobius hasta* | Bait primers^a^ |
| --- | --- | --- | --- |
| T1-F  -R | TATACACTACAGTTACTTGCCCG  GCCGATTTTGCCCACTACTA | AATCTTTCACCCGCTTGCC  TGAGCCCACTATTTACCCTTCA | AATCTTTCACCCGCTTGCC  TGAGCCCACTATTTACCCTTCA |
| T2-F  -R | GTTTCCCAAGTTCAACGGC  GGGTTTATCTCCGCTTCTGC | CAACATAGTATAAGAGGTCCTGCC  TGGGGTCTTCTCGTCTTATGT | **GGTATCCTGACCGTGCAAAG^b^**  **TGTAATTATCCCCGCTTCTGC** |
| T3-F  -R | CGCAATCCCCTTTTAGAGCC  CTCTTCTTTTCGGTCCTTTCGT | CTGGGATAACAGCGCAATCC  CCCTCTTTTCGGTCCTTTCG | TAGAGCCCATATCGACAAGAGG  CTTTTCGGTCCTTTCGTACTAGA |
| T4-F  -R | CCTTCCGCCTCCTTAGAAAG  CACGTTCTTGGGGTATGG | CTCCCGCCTCCTTAGAAAGA  CCAGTGCCTAGTACAATACCG | CTCCCGCCTCCTTAGAAAGA  CCAGTGCCTAGTACAATACCG |
| T5-F  -R | GCAGGACACTAACCCACATC  GGGGAGTAGTCGGATGCTC | CTGAATAAGATTGACAGGAACACT  GCAGGCGGGGTAAGAGT | TTGACAGGACACTAACCCGC  TCCCAAGTAGCAGGTTACCG |
| T6-F  -R | CACTTGATACTCAGTCATTTTAC  TGGTATTACTATAAAGAAAATTATTACA | ACAATCCACCACTTATCTCTCAG  GGTCATCCCCTAAAAGAGCC | GGGCTACAATCCACCACTTATC  ATCCCCTAAAAGAGCCCCAG |
| T7-F  -R | CATGCTTTTGTAATAATTTTCTTTATAG  TGGGGGGTAGACTGTTCA | ACCAGATTAACAATGTCATCGTG  CCAGAGGATGCAAGTAGAAGT | TGGGGCTCTTTTAGGGGAT  GCCAGAGGATGCAAGTAGAAG |
| T8-F  -R | CTCCCTGTCCTTGCTGC  AGGTTCTTTTTTGCCAGCAT | CTTCTTCTTCTATCACTCCCTGT  AGTAGGCGACAATATGCGAA | GTGCTTCTTCTTCTATCACTCCC  AGTAGGCGACAATATGCGAA |
| T9-F  -R | TGGGCATAGTCTGAGCCATA  GAGCCTCCATGCAGTGTAG | AGTTTGAGCAATGATGGCCA  TGATCATAGTAGGGGGGTTTCTC | GGGTACATGGGTATAGTTTGAGC  AATTGAGCCTCCATGCAGAG |
| T10-F  -R | CCAATAGTTGCAGTTATTACATC  ATTAGAGATACTGAATTAAAATGAAATC | TCTTACATCAGTAGTGCTCCTG  AAGTTATTAAGAGCGCGGCA | CCTGCTCCTTACCATCTTAGAGA  GGGTGGTTGTGTTGTAATGGA |
| T11-F  -R | CCTTGTTTTCCCCCAACTAC  CATGAGTACAAGAATAGATGCAAG | CTTGTATCTCTAGCTCTGTTCGC  TACCAAAATAGAGGCAAGTAGGG | CTTGTATCTCTAGCTCTGTTCGC  CCAAAATAGAGGCAAGTAGGGC |

^a^ A T7 promoter sequence, “TAATACGACTCACTATAGGG” was add to the 5’ end of the primers.

^b^ The primer pairs also used for qPCR experiment.
